# High-throughput engineering of cytoplasmic- and nuclear-replicating large dsDNA viruses by CRISPR/Cas9

**DOI:** 10.1101/2022.06.13.495503

**Authors:** Alberto Domingo López-Muñoz, Alberto Rastrojo, Rocío Martín, Antonio Alcamí

## Abstract

The application of CRISPR/Cas9 to improve genome engineering efficiency of large dsDNA viruses has been extensively described, but a robust and versatile method for high-throughput generation of marker-free recombinants for a desire locus has not been reported yet. Cytoplasmic-replicating viruses use their own repair enzymes for homologous recombination, while nuclear-replicating viruses use the host repair machinery. This is translated into a wide range of Cas9-induced homologous recombination efficiency depending on the virus replication compartment and viral/host repair machinery characteristics and accessibility. However, the use of Cas9 as a selection agent to target parental virus genomes robustly improves the selection of desired recombinants across large dsDNA viruses. We used ectromelia virus (ECTV) and herpes simplex viruses (HSV) type 1 and 2, to optimize a CRISPR/Cas9 method that can be versatilely used for efficient genome editing and selection of both cytoplasmic- and nuclear-replicating viruses. We performed a genome-wide genetic variant analysis of mutations located at predicted off-target sequences for 20 different recombinants, showing off-target-free accuracy by deep-sequencing. Our results support this optimized method as an efficient, accurate and versatile approach to enhance the two critical factors of high-throughput viral genome engineering: generation and color-based selection of recombinants. This application of CRISPR/Cas9 reduces time and labor of screening of desired recombinants, allowing for high-throughput generation of large collections of mutant dsDNA viruses for a desire locus in less than two weeks.

**DATA SUMMARY:** Raw sequence reads are available at the European Bioinformatics Institute (EMBL-EBI) European Nucleotide Archive (ENA) as Bioproject ID PRJEB32151 and PRJEB32152.

Six supplementary figures, eleven supplementary tables and supplementary methods are available with the online version of this article. The authors confirm all supporting data, code and protocols have been provided within the article or through supplementary data files.

## INTRODUCTION

Efficient, versatile, and reliable genome engineering of large double-stranded DNA (dsDNA) viruses is instrumental for pathogenesis and virulence studies, as well as for vaccine vector and oncolytic virus generation. Genome engineering by CRISPR/Cas9 has been shown in the recent years as a mighty platform to generate recombinant dsDNA viruses, such as vaccinia virus (VACV), herpesviruses and African swine fever virus (ASFV) (1–3). The Cas9-guide RNA (gRNA) complex cleaves the viral DNA, producing a double-stranded break (DSB) which is repaired by mammalian nuclear enzymes. This DSB can be fixed by non-homologous end joining (NHEJ), a template-free mechanism which incorporates indels, or by homology-directed repair (HDR) when a donor template is provided (4). CRISPR/Cas9 significantly promotes homologous recombination in the presence of donor templates, phenomenon which spontaneously occurs at very low frequencies during viral replication (5, 6). This low efficiency requires the use of marker genes to facilitate selection and screening. However, marker-free recombinants demand laborious selection methods and time-consuming screening of virus isolates from a large background of parental virus during traditional genome editing. This may become particularly challenging when generating marker-free recombinants from a color-labeled parental virus, where visual identification of recombinants becomes more complicated. Even in relatively efficient scenarios such as for VACV and other poxviruses, the efficiency of spontaneous homologous recombination remains between 0.1-1%, while it drops to a range of 0.01-0.001% for other DNA viruses such as herpesviruses, which rely on the host repair machinery (5–8). Poxviruses and other dsDNA viruses, such as ASFV, replicate within the cytoplasm of infected cells, while herpesviruses do so inside the cell nucleus. This is a key factor to understand the different efficiency of HDR between cytoplasm- and nucleus-replicating viruses after a Cas9-induced DSB. While CRISPR/Cas9 can greatly facilitate HDR through the nuclear host repair machinery for nucleus-replicating viruses, it can lead to inefficient repair for cytoplasm-replicating viruses since these depend on their own viral repair enzymes (8). The application of CRISPR/Cas9 to improve genome editing of herpesviruses has been extensively reported (1, 9). Nonetheless, CRISPR/Cas9 genome engineering of VACV has been described as inefficient tool to increase rates of homologous recombination, while it was shown to improve it up to 4 logs for ASFV (2). This suggests that despite cytoplasmic replication, the viral repair machinery determines the efficiency for increasing rates of HDR after a Cas9-induced DSB.

The two key steps in the dsDNA virus genome engineering by CRISPR/Cas9 are the initial generation of recombinants and the selection of these from a background of parental virus. Initial recombination means to increase rates of HDR, but the utility of CRISPR/Cas9 as an efficient selection tool also lies in its ability to cut parental genomes leaving most of them unable to propagate (8, 10). While CRISPR/Cas9 targeting does not improve recombination frequency over spontaneous homologous recombination for VACV, it significantly increases rates of HDR for HSV and ASFV (2, 8, 9). Interestingly, CRISPR/Cas9 can be efficiently used as a selection agent of recombinants for all these large DNA viruses (8, 10). A robust, efficient, and accurate CRISPR/Cas9 method that maximizes the efficiency of both the initial generation and selection of recombinant steps, as well as that can be versatilely used for cytoplasmic- and nuclear-replicating viruses, will facilitate the genome editing of other large DNA viruses extensively studied in the field.

Here we report an optimized protocol, termed as the Two-Progenies method, for generating large collections of recombinants by CRISPR/Cas9 for cytoplasmic- and nuclear-replicating dsDNA viruses. We used ECTV and HSV type 1 and 2 as models of cytoplasmic- and nuclear-replicating viruses, respectively. Our method exploits both the generation and selection of recombinant steps to maximize efficiency and reduce time and labor of screening of desired recombinants, allowing for off-target-free high-throughput generation of large collections of mutant dsDNA viruses for a desire locus in less than two weeks.

## METHODS

### Cells and viruses

293T (ATCC CRL-3216), BSC-1 (ATCC CCL-26) and Vero cells (ATCC CCL-81) were cultured in DMEM with 5% (v/v) Fetal Bovine Serum (FBS), 2 mM L-glutamine, and antibiotics (75 μg/ml penicillin, 75 U/ml streptomycin and 25 μg/ml gentamycin). All mammalian cells were cultured at 37° C in a CO2 buffered cell incubator and regularly tested for mycoplasma contaminations by PCR with primers Myco_Fw and Myco_Rv (Table S1). The source of the ECTV Naval isolate (accession no. KJ563295) was previously described (11). HSV-1 strain SC16 (accession no. KX946970 (12)) and HSV-2 strain 333 (accession no. LS480640 (13)) were kindly provided by Dr. Helena Browne, University of Cambridge (UK).

Recombinant viruses HSV-1 *UL26-27* Red and HSV-1 Red ΔgG-eGFP have not been described, while HSV-2 *UL26-27* Red and HSV-2 Red ΔgG-eGFP have been previously described (14). Those four viruses were made by a simplified CRISPR/Cas9 protocol where the *US4* locus encoding for glycoprotein G (gG) was replaced by an eGFP cassette in HSV-1/2 Red ΔgG-eGFP (detailed in Supplementary Methods, Figs. S1, S2 and S3). For viral stocks production, two confluent P150-cm^2^ plates of BSC-1 (for ECTV) or Vero cells (for HSV) were infected at a multiplicity of infection (MOI) of 0.1 plaque forming units (PFU)/cell and the viral inoculum was removed after 2 hours post-infection (hpi). Cells and supernatant were collected when reached 90-100% of cytopathic effect and cellular debris was concentrated by centrifugation at 300 x *g* during 5 min. After discarding the supernatants, 0.5 ml of 10 mM Tris-HCl pH 8.8 was added to the pellet, freezing/thawing three times. The pellet was lysed with a 23-gauge needle and 1 ml syringe for thirty times, releasing the intracellular virus. After 10 min of centrifugation at 500 x *g*, the supernatant was collected. This was done another two times. All the procedures were performed at 4° C using pre-cooled instrumentation. The supernatants were combined, aliquoted, frozen, and titrated twice by serial dilutions in duplicate BSC-1 (for ECTV) or Vero cells (for HSV) monolayers with 2 x 10^5^ cells/well in 12-well plates seeded the day before. After 2 hpi, viral inoculum was removed, and cells were overlaid with 1 ml of semi-solid carboxymethyl cellulose (1.5% w/v) 2% FBS DMEM. Cells were fixed in 10% formaldehyde at 48 hpi and plaques were stained with 0.1% (w/v) crystal violet for virus plaque counting.

### Plasmids

All sequence references correspond to the ECTV isolate Naval complete genome (KJ563295), HSV-1 strain SC16 partial genome (KX946970) and HSV-2 strain 333 complete genome (LS480640). All primers sequences can be found in Table S1.

To make the donor vector pF_195P-eGFP, two regions of the ECTV *195P* pseudogene were amplified from the ECTV Naval genome by PCR. The first region (187,133 – 187,461) was amplified with primers F_up(195P)_Fw and F_up(195P)_Rv, while the second region (187,574 – 187,890) with primers F_dw(195P)_Fw and F_dw(195P)_Rv. The VACV p7.5 promoter-eGFP cassette (see Supplementary Methods) was amplified from pF_INS-GFP (a kind gift of L. J. Sigal, Thomas Jefferson University), with primers p7.5-GFP_Fw and p7.5-GFP_Rv. Then, fragments were cloned into a modified version of pcDNA3.1/Zeo(-) (Invitrogen) by In-Fusion cloning (Takara Bio Inc.), obtaining pF_195P-eGFP. The modified pcDNA3.1/Zeo(-) was generated by PCR amplification, excluding the human cytomegalovirus (CMV) enhancer/promoter – bovine growth hormone polyadenylation signal (bGH) region with primers pcDNA3.1_Fw and pcDNA3.1_Rv.

The *US4* locus from HSV-1 SC16 genome (136,628-138,344) was amplified with primers F_up_US4(1)_Fw and F_dw_US4(1)_Rv and then, In-Fusion cloned into the modified pcDNA3.1/Zeo(-), obtaining plasmid pF_US4(1)_FL (Fig. S4a). Similarly, the *US4* locus from HSV-2 333 genome (137,411-140,507) was amplified with primers F_up_US4(2)_Fw and F_dw_US4(2)_Rv and In-Fusion cloned, to produce plasmid pF_US4(2)_FL (Fig. S4a). Each donor plasmid for the *US4* locus was generated by using pF_US4_FL as template (see Fig. S4a and Supplementary Methods). Donor plasmids pF_US4(1)/(2)_FL were linearized by PCR with primers pF_US4(1)_KO_Fw and pF_US4(1)_KO_Rv, and primers pF_US4(2)_KO_Fw and pF_US4(2)_KO_Rv, respectively, and In-Fusion circularized to generate pF_US4(1)/(2)_KO (Knock-Out) vectors, where the *US4* open reading frame (ORF) was excluded. Vectors pF_US4(1)/(2)_SC (Stop-Codon) were produced with primers pF_US4(1)_SC_Fw and pF_US4(1)_SC_Rv, and primers pF_US4(2)_SC_Fw and pF_US4(2)_SC_Rv, respectively, containing a non-translatable version of the *US4* gene. Plasmids pF_US4(1)/(2)_Δcyt (cytoplasmic tail), with primers pF_US4(1)_Δcyt_Fw and pF_US4(1)_Δcyt_Rv, and primers pF_US4(2)_Δcyt_Fw and pF_US4(2)_Δcyt_Rv, respectively, where gG cytoplasmic tail was excluded. Plasmid pF_US4(2)_mgG was made using primers pF_US4(2)_mgG_Fw and pF_US4(2)_mgG_Rv, excluding the secreted gG2 (sgG2) domain (Fig. S2a). Plasmid pF_US4(1)_gG2 was produced by amplification of gG2 ORF from pF_US4(2)_FL with primers pF_US4(1)_gG2_Fw and pF_US4(1)_gG2_Rv, and then, gG2 ORF was In-Fusion cloned into the linearized pF_US4(1) backbone, with primers F_dw_US4(1)_Fw and F_up_US4(1)_Rv. Similarly, gG1 ORF from pF_US4(1)_FL was amplified with primers pF_US4(2)_gG1_Fw and pF_US4(2)_gG1_Rv and In-Fusion cloned into previously linearized pF_US4(2) backbone, with primers F_dw_US4(2)_Fw and F_up_US4(2)_Rv, generating donor vector pF_US4(2)_gG1. The region of interest in every plasmid was fully Sanger sequenced with primers pZ_Fw and pZ_Rv.

### gRNA design and cloning

gRNAs were designed with Protospacer Workbench software (15). gRNAs designed to target the *195P* pseudogene of ECTV were based on region of interest (ROI) 187,462-187,573 (Table S2). gRNAs targeting the selected region of the CMV and bGH sequences (235-310 and 1917-2029, respectively in pEGFP-N3, Takara), were designed accordantly to each ROI (Table S3 and Supplementary Methods). Every gRNA was filtered using the Cas-offinder algorithm (including in the Protospacer Workbench software), allowing up to 5 mismatches. gRNAs with 0 off-targets were selected based on their Cas9 cleavage activity, defined by the Doench-Root Activity score (16), and location across their ROI. Each gRNA was generated by annealing of two complementary oligonucleotides, and then, the resultant dsDNA fragment was golden gate cloned in the *BbsI* sites of pSpCas9(BB)-2A-Puro V2.0 (Addgene #62988) or pSpCas9ΔNLS(BB)-2A-Puro plasmids. The latter was in-house generated by removing the nuclear localization signals from pSpCas9(BB)-2A-Puro V2.0 with four primers (pCas9ΔNLS_1 to 4) by In-Fusion cloning. gRNA_195P-2 was constructed with primers gRNA_195P-2_Fw and gRNA_195P-2_Rv, while gRNA_195P-4 with primers gRNA_195P-4_Fw and gRNA_195P-4_Rv. Then, they were cloned separately into plasmid pSpCas9ΔNLS(BB)-2A-Puro. gRNA_CMV-3 with primers gRNA_CMV-3_Fw and gRNA_CMV-3_Rv, giving pSpCas9(gRNA_CMV-3) and finally, gRNA_bGH-4 with primers gRNA_bGH-4_Fw and gRNA_bGH-4_Rv, to produce pSpCas9(gRNA_ bGH-4). The inserted gRNA cloned into every plasmid was fully Sanger sequenced with primer hU6_Fw.

### Generation of recombinant viruses by CRISPR/Cas9

Recombinant viruses were produced by following the same general workflow represented in Fig. S5, termed as the Two-Progenies method. In brief, 293T cell monolayers with 3 x 10^5^ cells/well in 6-well plates, were transfected using FuGENE HD (Promega) with 1 μg of the corresponding first pSpCas9-gRNA plasmid (for HSV; pSpCas9ΔNLS-gRNA for ECTV) to target the desired ROI, and 2 μg of the indicated “pF” donor plasmid (i.e., a mass ratio of 1: 2 for pSpCas9-gRNA: pF). After 8 h post-transfection, transfected cells were selected in the presence of 10 μg/ml of puromycin (Sigma). Then, after overnight treatment, selected cells were infected with indicated virus at a MOI of 0.1 PFU/cell, removing the viral inoculum 2 h later, washing, and adding 2 ml of fresh 10% FBS DMEM with puromycin. Both cell-associated and supernatant virus (first progeny) were harvested together after 48-72 hpi and subjected to three cycles of freezing-thawing. Simultaneously, 293T cell monolayers were transfected with 3 μg of the corresponding second pSpCas9-gRNA (or pSpCas9ΔNLS-gRNA) plasmid to target the same desired ROI. Transfected, puromycin-selected cells were infected with the clarified supernatant from the first progeny, being removed after 2 h, washed, and overlaying with 2 ml of fresh 10% FBS DMEM with puromycin. Virus present in both cells and supernatants (second progeny) was harvested after 48-72 hpi, and freeze-thawed three times. The second progeny was serially diluted and used to infect Vero (for HSV) or BSC-1 (for ECTV) cell monolayers in 12-well plates, adding semisolid carboxymethyl cellulose (1.5% w/v) 2% FBS DMEM after inoculum removal. Isolated recombinant viral plaques were identified 48-96 hpi by fluorescent microscopy in a Leica DM IL LED inverted microscope equipped with a Leica DFC3000-G digital camera, also used to take micrographs. Five rounds of plaque isolation, PCR screening, SDS-page, Sanger and deep sequencing were performed to confirm the desired genomic modification for each case. For PCR screening, primers E194_Fw and 5GFP_Rv for ECTV, while primers Up_US4(1)_Fw and Dw_US4(1)_Rv for HSV-1, and primers Up_US4(2)_Fw and Dw_US4(2)_Rv for HSV-2 were used.

### Deep sequencing

Viral stocks were treated with DNAse I (0.25 U/μl, Roche), RNAse A (20 μg/ml, Roche) and nuclease S7 (0.25 U/μl, Roche) and then incubated 2 h at 37° C to eliminate cellular DNA/RNA. Sterile EDTA (12 mM) and EGTA (2 mM) at pH 8 were added to inactivate nuclease activity. Then, viral particles were lysed by incubation with Proteinase K (200 μg/ml) and sodium dodecyl sulfate (0.5%) for 1.5 h at 45° C. Viral DNA was purified by Phenol:Chloroform:Isoamyl Alcohol (25:24:1, v/v) extraction. All the procedures were performed under sterile conditions. DNA was quantified in a Nanodrop One spectrophotometer (Thermo Fisher Scientific) and by fluorometric quantitation using a PicoGreen device (Invitrogen). Contaminating DNA was checked by PCR against mycoplasma, prokaryotic 16S rRNA (primers 16S_Fw and 16S_Rv) and eukaryotic 18S rRNA (primers 18S_Fw and 18S_Rv) (17, 18). Finally, viral DNA was tested to determine HSV type cross-contamination by PCR (Up_US4(1)_Fw and Dw_US4(1)_Rv for HSV-1; Up_US4(2)_Fw and Dw_US4(2)_Rv for HSV-2). An aliquot of viral genomic DNA (100 ng) was submitted to MicrobesNG, University of Birmingham (UK) for sequencing. Illumina libraries were prepared with NEBNext Ultra DNA Library Prep Kit (New England Biolabs). DNA samples were fragmented in a Covaris instrument and sequenced on an Illumina MiSeq device as paired-end reads, 2 x 250 base pairs (bp), according to manufacturer’s recommendations. Sequencing statistics for every sample used in this study can be found in Table S4.

### Genome-wide analysis of genetic variants at predicted Cas9 off-target sequences

Reads from each sequenced sample were trimmed using Trimmomatic v0.36 (19), quality-filtered with PrinSeq v1.2 (20), and aligned against the indicated parental genomic sequence for each case, by using Bowtie 2 v2.3.4.1 with default settings (21). Bowtie 2 alignments were visualized and screened using Integrative Genomics Viewer v2.8.2 (22) to detect large gaps and rearrangements. Minor variants (MVs) were detected as previously described (23). Briefly, VarScan v2.4.3 (24) was used with settings intended to minimize sequencing-induced errors from the raw calling of MVs. Detected MVs from VarScan calling were then annotated onto their corresponding genome, to determine their mutational effects. MVs were then additionally filtered by coverage > 200. An additional filter was implemented to detect those MVs with high frequency but low coverage, where read depth at the given position had to be greater than the product obtained from dividing 200 (coverage threshold) by the variant frequency (0–100) at the given position (23). MVs were considered as *de novo* appearance when, after coverage filtering, its frequency in the previous parental viral population was < 0.01. *De novo* MVs were examined to determine whether they were located within any of the predicted Cas9 off-target sequences for both gRNAs (gRNA_CMV-#3 and gRNA_bGH-#4) used to generate each recombinant. For full lists of MVs detected in each viral population, see Table S5 (parental and recombinant HSVs-1), Table S6 (parental and recombinant HSVs-2) and Table S7 (recombinant ECTV 195P-eGFP). For a summary list of detected, filtered, and categorized MVs for every sample sequenced in this study, see Table S8.

### Statistical analysis

Data analyses were performed using GraphPad Prism 8 (v8.4.3) software. Student’s unpaired *t-* test or two tailed Mann-Whitney *U*–test were used. Differences in both tests were considered statistically significant when *p*-values were below 0.05.

## RESULTS

### Optimized CRISPR/Cas9 protocol for engineering cytosol- and nuclear-replicating dsDNA viruses

We have developed a versatile, fast, and reliable workflow for scientific and therapeutic applications involving large dsDNA viruses. Our method comprises two steps, initial recombination followed by one round of Cas9 selection (Figs. 1a, 1b and S5). First, a Cas9-mediated HDR targets the locus of interest with one gRNA (first viral progeny), then one round of Cas9 selection targets again the locus of interest at a different and intact gRNA site (second viral progeny), taking in total less than one week (virus replication cycle dependent). We adapted our method for engineering of viruses with cytoplasmatic replication, such as ETCV, by removing the nuclear localization signals (NLS) when transiently expressing Cas9 (25). For viruses with nuclear replication (i.e., herpesviruses), Cas9 is expressed containing NLS. For initial validation, we selected the *195P* locus within the ETCV genome since this is a pseudogene which disruption does not affect the viral phenotype (11). We also selected the eGFP cassette within the genome of previously generated recombinant HSV-1/2 (Figs. S2 and S3).

**Fig. 1.**
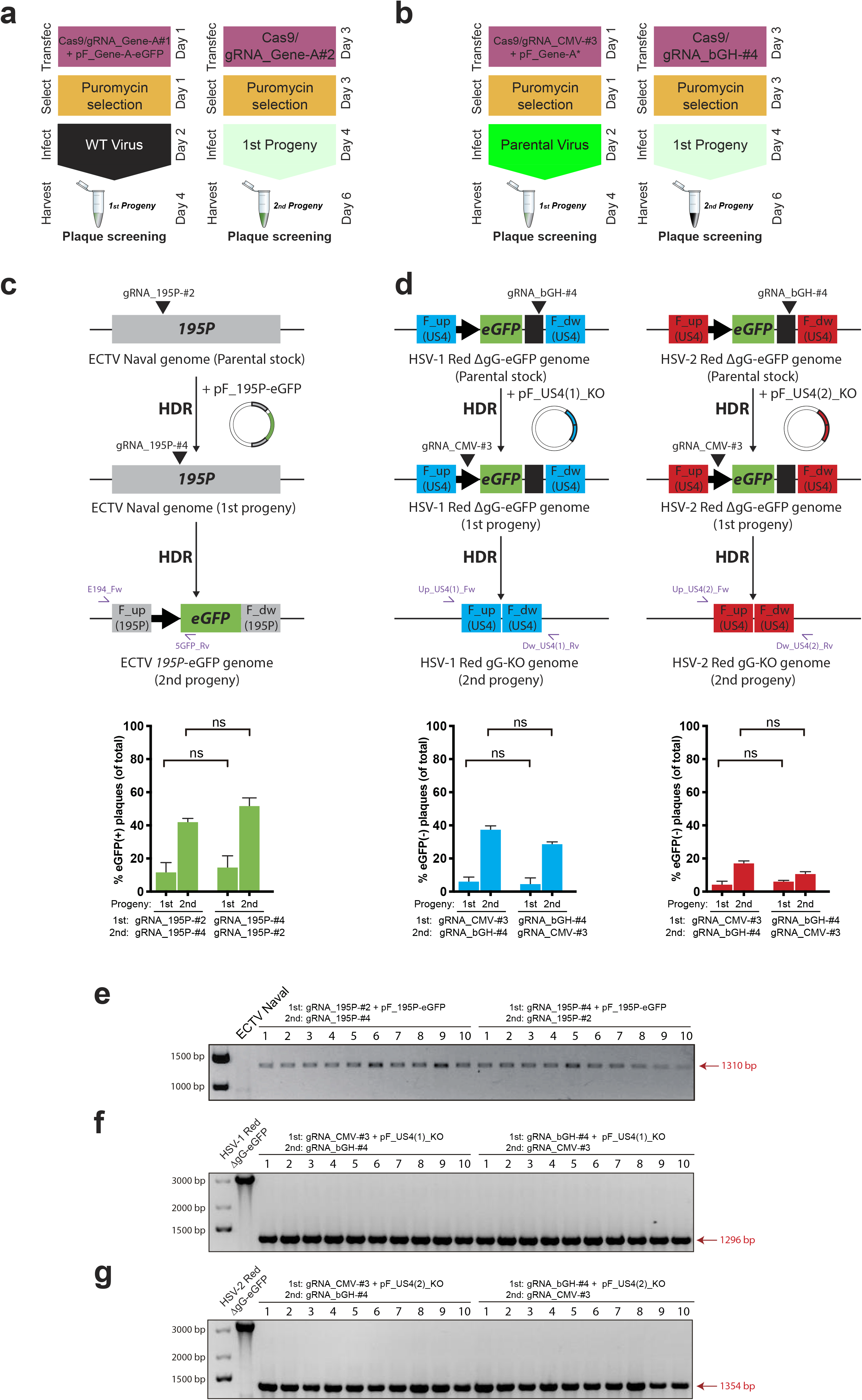
Efficient and rapid generation of CRISPR/Cas9-derived recombinant ECTV and HSV-1/2 using the Two-Progenies method. (**a, b**) Schematic workflow showing the color-based protocol to create knock-in (**a**) and marker-free (**b**) recombinants for a desire locus (e.g., Gene *A*). (**c, d**) Schematic of parental and resulting recombinant genomes for ECTV (**c**), HSV-1 (**d**, center) and HSV-2 (**d**, right), as well as plasmids for HDR, gRNA target sites, and primers for PCR verification (purple arrows) used for each progeny generation. (**c**) An eGFP cassette is inserted into the *195P* locus of the ECTV genome. (**d**) An eGFP cassette already present in the parental recombinants HSV-1/2 Red ΔgG-eGFP is completely removed. Second progenies from ECTV (**c**), HSV-1 (**d**, center) and HSV-2 infections (**d**, right) were collected, serially diluted, grown on fresh cell monolayers, and scored for green fluorescence by microscopy. Plaques in the lowest countable dilution per condition are graphed showing mean and SEM of two independent experiments (*ns* = not significant, *p* > 0.05 by Student’s unpaired *t*-test). Experiments were performed with reversed order of each pair of gRNAs used to target the same locus. (**e, g**) PCR verification of the desired sequence of selected recombinants. Ten random viral plaques from each second progeny and each pair of gRNAs combination were hand-picked, isolated, and PCR-tested. (**e**) PCR screening of 20 recombinant eGFP(+) ECTV *195P*-eGFP plaques, with the wildtype virus serving as the negative control (ECTV Naval). (**f**) PCR screening of 20 eGFP(-) HSV-1 Red gG-KO plaques, with the parental recombinant virus serving as control (HSV-1 Red ΔgG-eGFP). (**g**) PCR screening of 20 eGFP(-) HSV-2 Red gG-KO plaques, with the parental recombinant virus serving as control (HSV-2 Red ΔgG-eGFP).

To generate the first viral progeny, we targeted the ETCV genome with gRNA_195P-#2 and induced HDR by proving a donor template containing eGFP (Fig. 1c). To do so for HSV-1/2 Red ΔgG-eGFP, we targeted the bGH site within the eGFP cassette with gRNA_bGH-#4 including a donor template to remove the eGFP cassette by HDR (Fig. 1d). The second recombinant progeny was then subjected to one round of selection by Cas9 targeting again the *195P* locus with gRNA_195P-#4 for ECTV (Fig. 1c), and the eGFP cassette with gRNA_CMV-#3 for HSV-1/2 (Fig. 1d). This round of Cas9 selection applied selective pressure over wild-type (WT) genomes in favor of the already recombined ones. After initial recombination, eGFP was detected in approx. 15-18% of the first progeny plaques for ECTV, independently of whether those were generated either with gRNA_195P-#2 or gRNA_195P-#4 (Fig. 1c bottom). Loss of eGFP signal for recombinant HSV-1/2 was detected in 8-10% of the first progeny plaques for both viruses, without differences between gRNA_bGH-#4 and gRNA_CMV-#3 (Fig. 1d bottom). Second progenies were also screened for eGFP signal after Cas9 selection was applied. Between 45-55% of the screened plaques were eGFP(+) for ECTV (Fig. 1c bottom). Negative plaques for eGFP reached a 35-40% of the recombinant HSV-1 second progeny plaques, while a more discrete 15-20% was observed for recombinant plaques of HSV-2 (Fig. 1d bottom). Interestingly, we found no significant differences in the efficiency of recombinant virus generation based on the gRNA combination, for any of the three viruses analyzed. Finally, we randomly hand-picked, isolated and PCR-checked 20 recombinant second progeny plaques from each virus (eGFP(+) plaques for ECTV, eGFP(-) for HSV), 10 from each gRNA combination. Of the 10 plaques from each virus and gRNA combination, all had the desired recombinant sequence (Fig. 1e-g). These data evidenced that our Cas9-based protocol allows efficient generation and selection of dsDNA viruses in less than one week, where the order of usage for a certain pair of gRNAs does not alter the intrinsic cumulative efficiency of those gRNA to generate and select the desired recombinant virus.

### The Two-Progenies method applies selective pressure to parental and NHEJ-ed genomes and efficiently selects recombinant dsDNA viruses

The two critical obstacles associated with dsDNA virus genome engineering are the initial generation of recombinants and the selection of these from a pool of parental virus. The identification and isolation of mutants can become extremely challenging and laborious when generating marker-free colorless recombinants, particularly within a large background of fluorescent parental virus. We observed that CRISPR/Cas9 improved the production of initial recombinants by enhancing HDR with the first gRNA (Fig. 1c, 1d). Cas9 inhibits replication of targeted unrepaired parental genomes, favoring the expansion of already recombined genomes where the gRNA target site has been lost as a result of the recombination event (8, 10). However, undesired NHEJ-ed recombinants, which may have lost the gRNA target site, are also able to escape a Cas9 selection driven by the same gRNA used for the initial recombination (Fig. 2a) (10). Since random InDels generated by NHEJ are usually small (1-10 bp) but can reach up to 200 bp (26, 27), we designed both gRNAs targeting the eGFP cassette in HSV-1/2 Red ΔgG-eGFP as far as possible from the eGFP ORF, i.e., at the edges of the CMV and bGH regions (Fig. 2a, 2b). We confirmed that a transfected plasmid encoding for eGFP in the absence of both CMV and bGH (a modified version of the pF_US4(1)/(2)_ΔgG-eGFP, see Fig. S1b) was sufficient for fluorescent detection. By doing this, we enhanced the selection step of our protocol towards HDR-ed genomes, by using the second gRNA to target an intact site only directed to parental and NHEJ-ed genomes during the second progenies generation (Fig. 2b). Remarkably, due to the fact that the parental HSV-1/2 Red ΔgG-eGFP viruses conserved the expression of the red marker (inserted in a different locus, see Fig. S2 and Supplementary Methods), we were able to observe in the pre-harvested second progenies shown in Fig. 1d, red infected cells which had lost the eGFP signal (Fig. 2c). This indicated that those cells contained mainly if not only HDR-ed recombinant viruses. As shown in Fig. 1e-g, none of the recombinant plaques randomly selected showed evidence of editing by NHEJ, i.e., a false positive. This suggested that our strategy inhibits both parental and NHEJ-ed genomes while allowing only HDR genomes, so desired recombinant, to replicate and increase their frequency in the second progeny. Altogether, these data evidenced that our Cas9-based protocol allows efficient, reliable generation and sensitive selection of dsDNA viruses in less than one week, where the frequency of desired recombinants after just one selection step is high enough to easily identify color-based recombinant plaques by fluorescent microscopy.

**Fig. 2.**
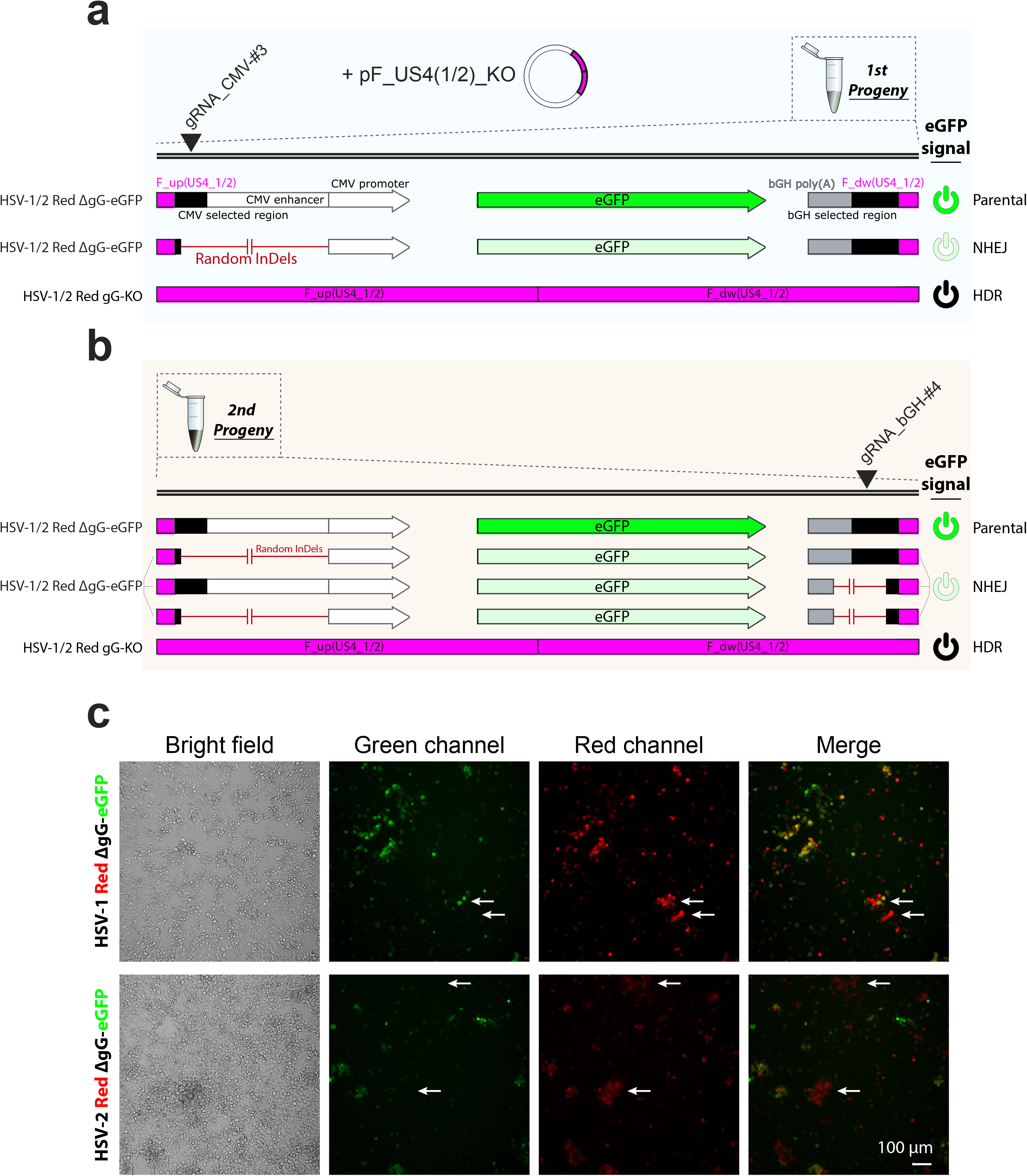
The Two-Progenies method applies selective pressure to the parental virus and can be efficiently used for visual selection of colorless recombinant dsDNA viruses. (**a**) Schematic of parental and theoretical resulting recombinant genomes obtained after the generation of the first viral progeny (e.g., HSV-1/2 Red gG-KO). This progeny may contain a mixture of parental (eGFP^High^), NHEJ-ed (eGFP^Low^) and HDR-ed (eGFP(-)) recombinants. Random InDels at the CMV targeted site, incorporated by NHEJ, does not disrupt eGFP expression. (**b**) Schematic of resulting recombinant genomes obtained after the generation of the second progeny (e.g., HSV-1/2 Red gG-KO). This final progeny may also include parental (eGFP^High^), NHEJ-ed (eGFP^Low^) and HDR-ed (eGFP(-)) recombinants. Random InDels in both *CMV* and *bGH* targeted sites do not abrogate eGFP expression. (**c**) Fluorescence micrographs of second progenies from HSV-1/2 Red gG-KO cultures after 48 hpi with the corresponding first progeny, following the above workflow. Evident cytopathic effect revealed infected cell whose viruses had lost eGFP expression but preserved Red signal (white arrows).

### The Two-Progenies protocol can be efficiently used for high-throughput generation of large collections of mutant dsDNA viruses for a desire locus

To further characterize the efficiency and accuracy of our color-based method to minimize the selection and screening of false negative recombinants, we used the same fluorescent HSV-1/2 Red ΔgG-eGFP model viruses as in the last figures, replacing the *eGFP* cassette with different donor templates containing modified versions of the *US4* locus (Figs. 3a and S4a). This allowed for a more comprehensive characterization of the efficiency of initial recombination (first progeny) and Cas9 selection steps (second progeny) when varying the donor template responsible for triggering HDR. We detected eGFP(-) recombinant plaques by fluorescence microscopy at a frequency ranging from 2% to 8% for HSV-1 and from 3% to 10% for HSV-2, donor-template-dependent, in the plated first progenies (Fig. 3b). The frequency of eGFP(-) recombinants increased in every condition and both HSV, ranging from 10% up to 70% when second progenies were plated for visual screening (Fig. 3b). Compiling the frequencies of eGFP(-) plaques by progenies within each HSV (including gG-KO mutants from condition 1^st^ gRNA_CMV-#3, 2^nd^ gRNA_bGH-#4 in Fig. 1d), we observed a significant increment of desired recombinant viruses at an average frequency of approx. 20% for HSV-1 and 35% for HSV-2 after the Cas9 selection step in the second progenies (Fig. 3c). We also tested by PCR ten random eGFP(-) plaques from each second progeny in Fig. 3b, finding that all had the desired recombinant sequence (not shown). From each recombinant virus, we selected two viral clones out of ten for further characterization. Protein expression was tested by western blot (Fig. S4c), and absence of large genomic gaps and rearrangements was confirmed by deep sequencing. These results showed how the frequency of desired recombinants is reliable and high enough to identify these mutants directly by visual screening, after just one selection step with a different secondary gRNA, independently on the HDR template provided. Our protocol allows for a robust generation and fast visual identification of large collections of marker-free recombinant viruses for a desire locus, avoiding the traditional initial screening of tens of marker-free recombinants by PCR.

**Fig. 3.**
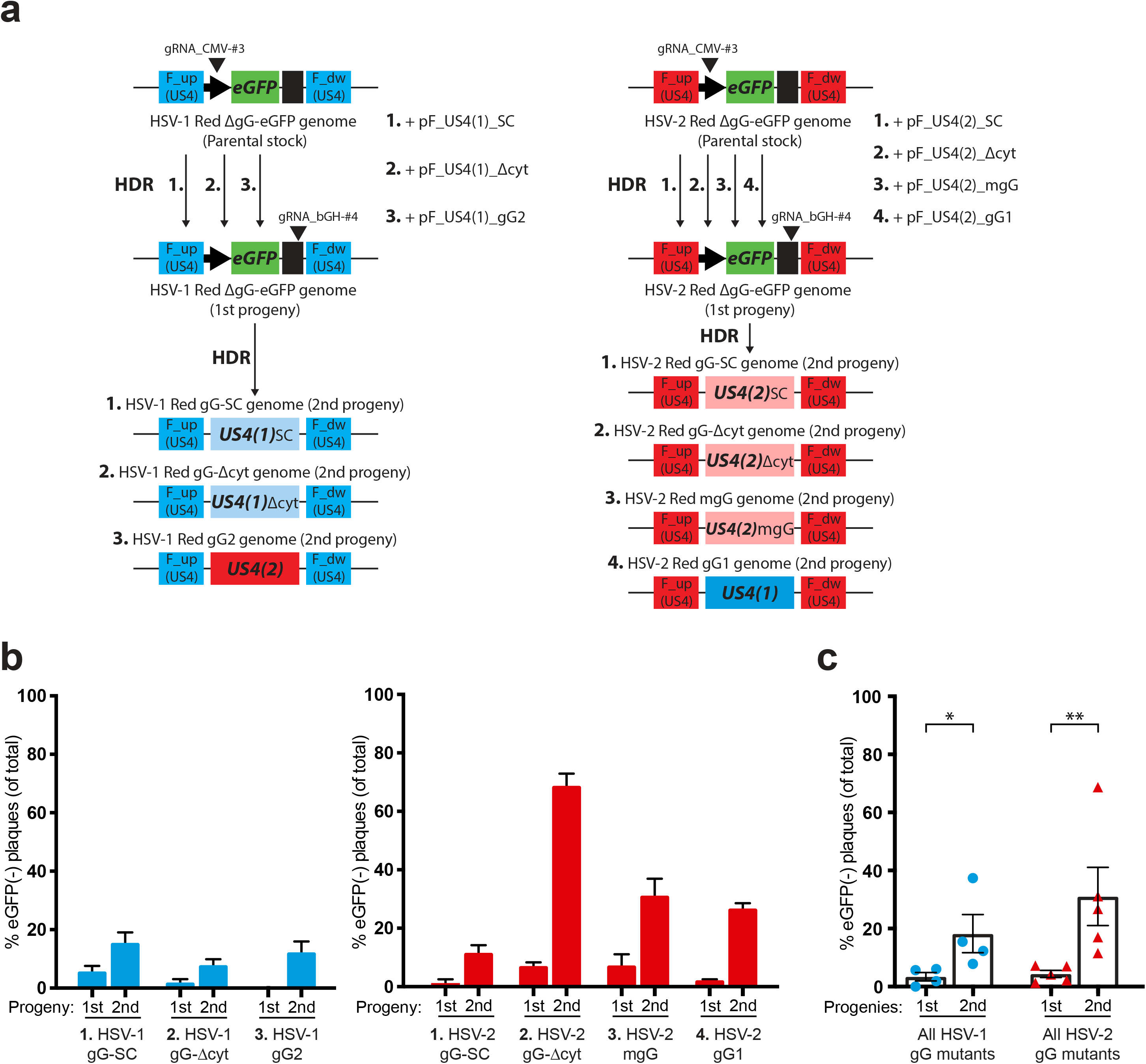
High-throughput generation of a marker-free collection of mutants HSVs-1/2 in the *US4* locus, using the Two-Progenies method. (**a**) Schematic of parental and resulting recombinant genomes for HSV-1 (left) and HSV-2 (right) in the *US4* locus, as well as donor plasmids for HDR and gRNA target sites. (**b**) First and second progenies from HSV-1 (left, blue bars) and HSV-2 infections (right, red bars) were collected, serially diluted, grown on fresh cell monolayers, and scored for green fluorescence by microscopy. Plaques in the lowest countable dilution per condition are graphed showing mean and SEM of two independent experiments. (**c**) Compiled frequencies of eGFP(-) plaques from first and second progenies in (**b**) are graphed showing mean and SEM (* *p* = 0.0286, ** *p* = 0.009 by two tailed Mann–Whitney *U*-test), including gG-KO mutants from **Fig. 1d** (condition 1^st^ gRNA_CMV-#3, 2^nd^ gRNA_bGH-#4). HSV-1 (left, blue circles) and HSV-2 infections (right, red triangles).

### The Cas9-based Two-Progenies protocol offers off-target-free accuracy confirmed by deep sequencing

The fact that large dsDNA viruses have a small genome (150-200 Kbp) compared to bacteria and eukaryotic organisms, minimizes the potential off-target sites for Cas9. However, the presence of complex repeated regions as well as other secondary structures within dsDNA viral genomes might be translated into a higher Cas9 promiscuity. Viral factors participating in the replication phase, together with an intrinsic high recombination rate exhibited by poxviruses and herpesviruses could also reduce Cas9 precision. Moreover, the fact that herpesviruses can generate a great degree of genetic diversity when replicating *in vitro* also calls into question the off-target-free accuracy of Cas9 when engineering dsDNA viruses (23, 28). In order to address this matter, we deep sequenced two isolates of every recombinant virus generated by the Two-Progenies method, as well as the parental viruses (Table S4). We did not observe large genomic gaps or rearrangements across the genome of any sequenced recombinant. We then performed a genome-wide genetic variant analysis to detect pre-existing and *de novo* appeared MVs within each recombinant isolate. We consistently detected a higher number of *de novo* MVs among HSV-2 recombinants than among HSV-1 counterparts (Fig. 4, Table S8). Nonetheless, we previously reported that HSV-2 evolves *in vitro* faster than HSV-1 (23). The fact that these recombinants were plaque-isolated five times in cell culture might explain the appearance of these *de novo* MVs. To determine if these *de novo* MVs had appeared due to *in vitro* genetic drift rather than being caused by Cas9-induced NHEJ events, we examined whether any of the detected *de novo* MVs were localized within any of all predicted Cas9 off-target sequences for both gRNAs used to generate these recombinants. Of the 20 sequenced recombinants analyzed, we were not able to find a single de novo MV located within a predicted off-target sequence with less than 10 mismatches (Table S9). Similar results were obtained when parental Cas9-derived HSV-1/2 Red ΔgG-eGFP were analyzed for *de novo* MVs located in predicted Cas9 off-target sequences for both gRNAs used (Fig. S6). Altogether, these results confirm that our method using Cas9 for high-throughput genome engineering of dsDNA viruses offers reliable genome-wide off-target-free accuracy.

**Fig. 4.**
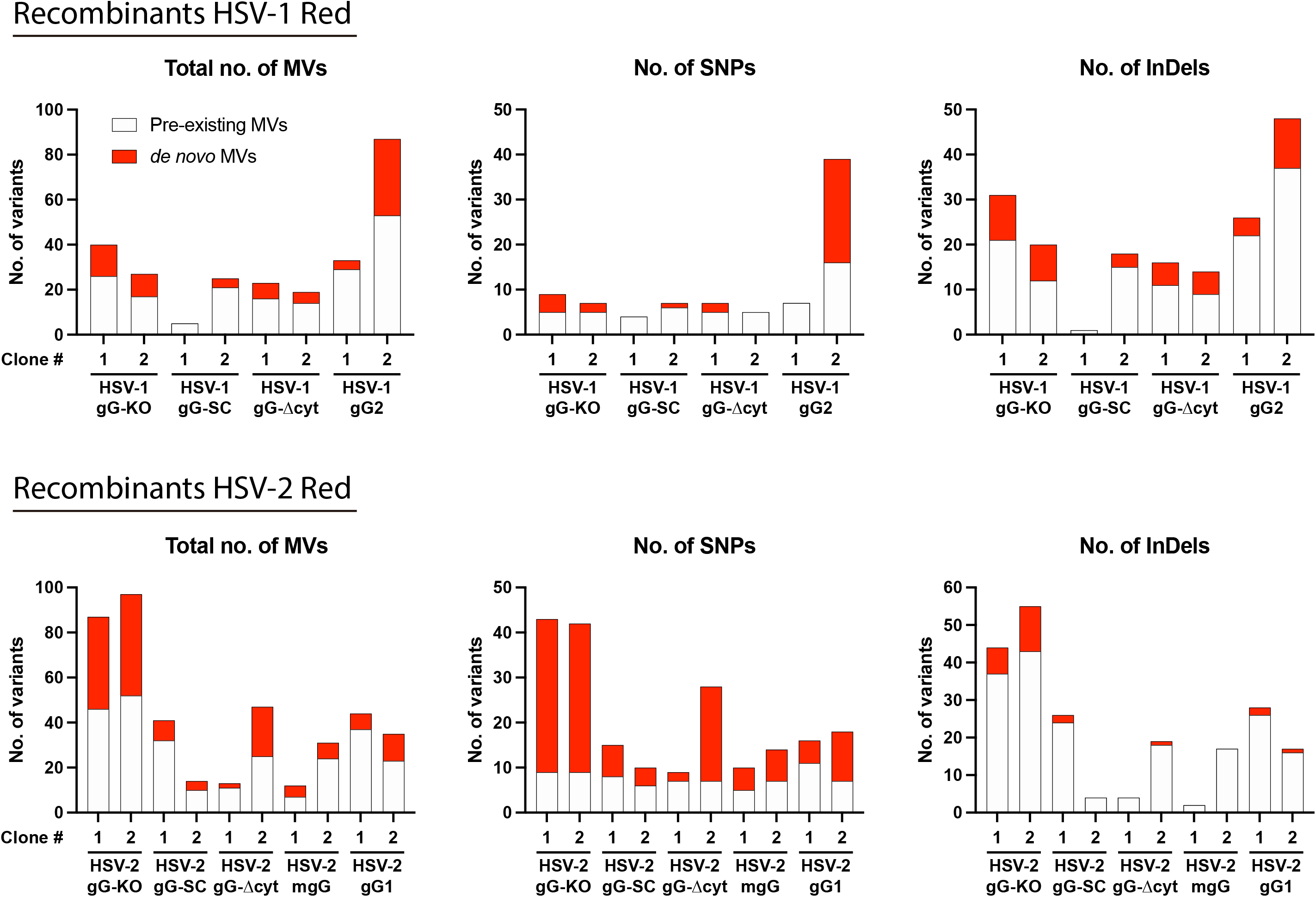
The Two-Progenies protocol offers off-target-free accuracy. Number of MVs, SNPs, and InDels are plotted according to genetic variant analysis data for each recombinant HSV. Histograms differentiate between pre-existing MVs observed in the corresponding parental virus (white) and *de novo* appeared MVs (red).

## DISCUSSION

The ability to quickly identify and select recombinant DNA viruses has wide and profound implications for viral pathogenesis and virulence studies, as well as for vaccine vector and oncolytic virus generation. CRISPR/Cas9 has proven to be groundbreaking for genome engineering of large dsDNA viruses. Several studies have evaluated the specificity and ability of this system to edit the HSV genome (1, 9). The first application to HSV reported an efficiency higher than 50% generating *US8* knock-out mutants by NHEJ, but decreasing bellow 10% for knock-in recombinants by HDR at the same locus (29). Other authors described HDR efficiencies of 8% and 23% when inserting or replacing fluorescent marker genes within the HSV genome (7, 30). Optimization refinements have been reported, describing highly variable HDR efficiency rates inserting or replacing different markers. Whereas a 48% of eGFP(+) plaques was obtained after the insertion of that marker gene in an intergenic region of the HSV genome, a dramatic reduction to 1% was found when replacing the viral gene *ICP0.* Then, substitution of eGFP by other marker genes, such as mCherry or LacZ genes, gave HDR rates of 10% and 2.1%, respectively (31). Nonhomologous insertion strategies, based on NHEJ, reaches 45% efficiency, but its fidelity varies (70-100%) based on genomic location (32). In contrast, we describe here a significant increment in HDR efficiency after one round of Cas9 selection for marker-free herpesviruses, reaching up to 70% in the best case. There have been also descriptions of CRISPR/Cas9 application to poxviruses, using VACV as the model poxvirus (25, 33). More recently, Gowripalan et al. concluded that Cas9-induced NHEJ or HDR is not efficient repairing the VACV genome (8), i.e., that CRISPR/Cas9 does not increase rates of homologous recombination in this poxvirus, but it is a highly efficient selection tool after initial spontaneous homologous recombination. Genome editing efficiency of poxviruses by spontaneous homologous recombination ranges from 0.1 to 1%. We report here for the first time that CRISPR/Cas9 can be used to increase Cas9-induced HDR up to 15-18% for ECTV, being also a highly efficient selection tool for selection of desired recombinants from a large background of parental virus. It is unknown to what extent genomes from each poxvirus are differentially exposed in the cytoplasm prior being contained into the viral factories, which might be an impediment to Cas9 access and targeting (34). Moreover, poxviruses are defined by their nuclear independence, which suggest that cleaved genomes of different poxvirus may not have the same access to host repair factors for NHEJ and HDR (35). On top of this, the exclusive properties of each poxvirus’ replication machinery may also differentially contribute to genome repair efficiency by HDR after Cas9 targeting. Other studies have also showed the advantage of using CRISPR/Cas9 to improve HDR efficiency for other cytoplasmic-replicating viruses, such as ASFV, which also establishes cytoplasmic viral factories for replication (2, 3). The potential of our method can be seen in the fact that we have optimized both critical steps of virus genome engineering, i.e., the production of initial recombinants (first progeny) and the selection of these from the background of parental virus (second progeny), independently on the virus replication compartment. Even in cases like VACV, where Cas9 targeting does not increase rates of HDR over spontaneous homologous recombination, the first progeny step will help to select early recombined genomes by Cas9 targeting with the first gRNA.

The generation of knock-out recombinants is often used as a research tool to study the contribution of a certain locus or gene to fundamental aspects of the virus biology. Once a relevant locus is identified, the detailed study of its mechanistic contribution to the virus life cycle requires the generation of additional recombinant viruses expressing variant forms of the protein of interest in order to analyze the role of protein motifs and domains played during infection. This implies a time-consuming, laborious, and expensive task when studying large multidomain viral proteins. It can get even more complicated if marker genes remain as a consequence of the genome engineering process, since the presence of those marker genes are typically unacceptable in therapeutics and vaccines. Our Two-Progenies method allows for high-throughput generation of large collections of mutant dsDNA viruses for a desire locus in less than 2 weeks (Fig. 5). By replacing the locus of interest by a marker gene, and then substituting it for each final desired mutant version of that viral gene object of study, we facilitate a visual, color-based identification and selection of desired recombinants at each step. This approach speeds up the generation of a large collection of recombinants since laborious and time-consuming rounds of plaque-picking, isolation and mass PCR screening are substituted for visual selection, relegating the traditional PCR screening to a confirmatory final step for already identified isolates as the desired recombinants. The fact that gRNAs targeting the eGFP cassette are designed far enough from the eGFP ORF and from each other, helps to avoid loss of fluorescence caused by NHEJ events, so it reduces the likelihood of selecting false positive recombinants. By using two different gRNAs, far away from each other to target a desired locus in two consecutive steps, parental and NHEJ-ed genomes suffer constant selective pressure due to Cas9 targeting, while HDR-ed recombinants replicate freely increasing their frequencies (Fig. 2). The fact that large dsDNA viral genomes are small when compared to bacterial and eukaryotic genomes reduces the potential number of sequences for Cas9 off-targeting. Because of this, Cas9-derived recombinants are not routinely screened for genomic rearrangements and MVs. This is particularly relevant for viruses which generate a high degree of genetic heterogenicity due to genetic drift in cell culture, such as HSV.

**Fig. 5.**
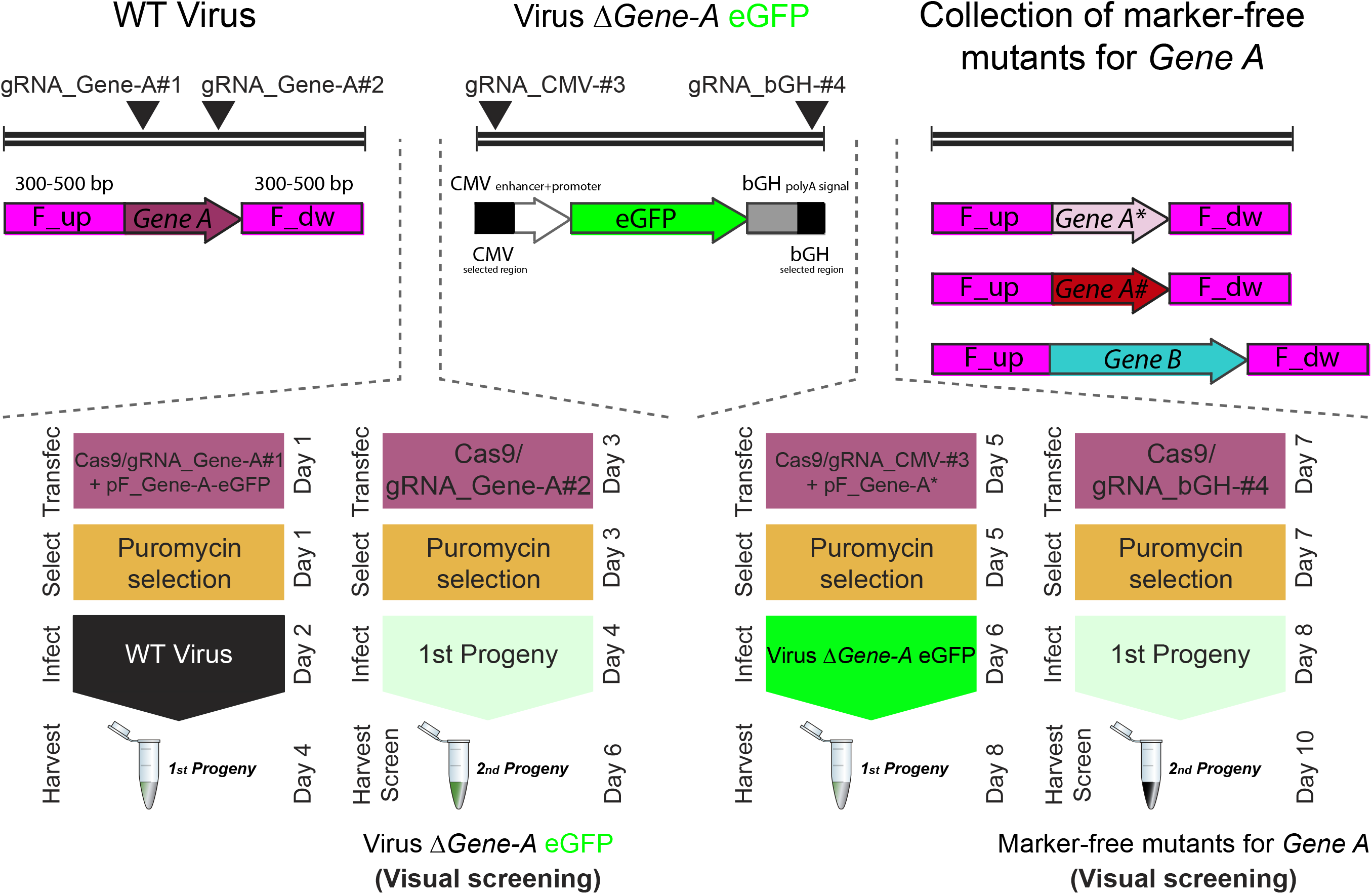
Schematic showing the color-based strategy to generate a large collection of Cas9-derived recombinant dsDNA viruses for a desire locus of interest (e.g., *Gene A*). The Two-Progenies protocol is used twice, first to replace the locus of interest by an eGFP cassette, and second, to substitute the latter by each mutant version of the locus of interest.

Altogether, our refinements contribute to improve and accelerate the accurate genome engineering of large dsDNA viruses, by increasing HDR efficiency using Cas9 as the initiator of recombination, and/or by using it as a selection agent to increase the frequency of desired recombinants over a large background of non-desired recombinant and parental viruses.

## Abbreviations

ECTV: ectromelia virus;
HSV: herpes simplex virus;
VACV: vaccinia virus;
ASFV: African swine fever virus;
gRNA: guide RNA;
DSB: double-stranded break;
NHEJ: non-homologous end joining;
HDR: homology-directed repair;
gG: glycoprotein G.

## AUTHOR STATEMENTS

### Authors and contributions

Conceptualization: ADLM, AR, AA; methodology: ADLM, AR; software: ADLM, AR; formal analysis: ADLM, AR; investigation: ADLM, AR, RM; resources: AA; data curation: ADLM, AR; writing - original draft preparation: ADLM; writing - review and editing: all authors; visualization: ADLM; supervision: AR, AA; funding: AA.

## Acknowledgements

We are grateful to the Genomics and Next Generation Sequencing Service at Centro de Biología Molecular Severo Ochoa for their support and advice. Genome sequencing was provided by MicrobesNG (http://www.microbesng.uk), which is supported by the BBSRC (grant no. BB/L024209/1).

