## Supplementary Material for "High-throughput engineering of cytoplasmic- and nuclear-replicating large dsDNA viruses by CRISPR/Cas9"

**This PDF file includes:**

Supplementary Figs. S1 to S6

Supplementary Methods

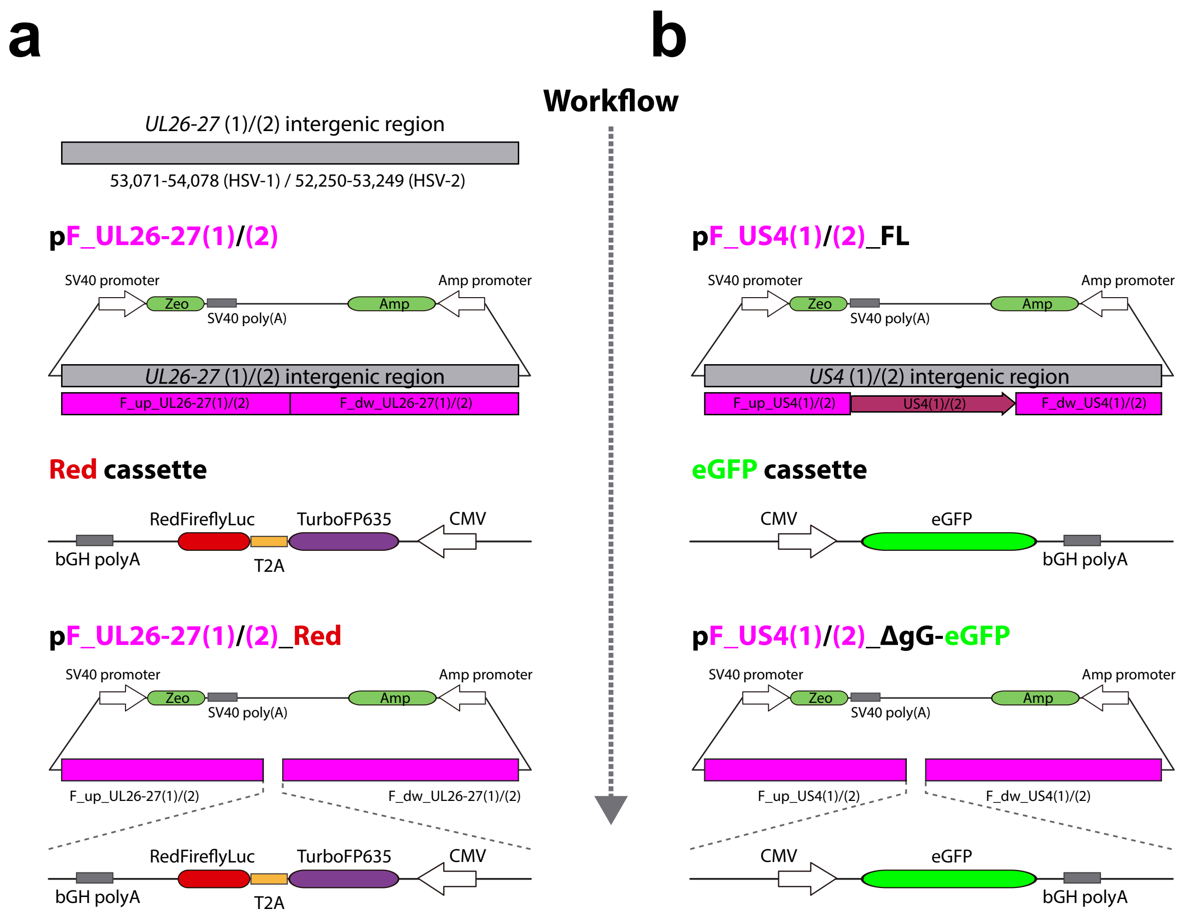


**Fig. S1. Schematic of the donor plasmids used as HDR templates for the generation of (a) HSV-1/2 *UL26-27* Red and (b) HSV-1/2 Red ΔgG-eGFP.** (**a**) Donor plasmids pF_UL26-27(1/2) were generated by PCR amplification of the *UL26-27* intergenic region from HSV-1/2 genomes and then, cloned into a modified pcDNA3.1 vector. A Red cassette (CMV-TurboFP635-T2A-RedFireflyLuc-bGH) was reversely cloned into the linearized pF_UL26-27(1/2), obtaining pF_UL26-27(1/2)_Red plasmids. (**b**) Donor plasmids pF_US4(1/2)_FL were constructed by amplification of the *US4* genic region from viral genomes and cloned into the modified pcDNA3.1 vector. An eGFP cassette (CMV-eGFP-bGH) was cloned into the linearized pF_US4(1/2)_FL, excluding the *US4* CDS, and producing pF_US4(1/2)_ΔgG-eGFP.

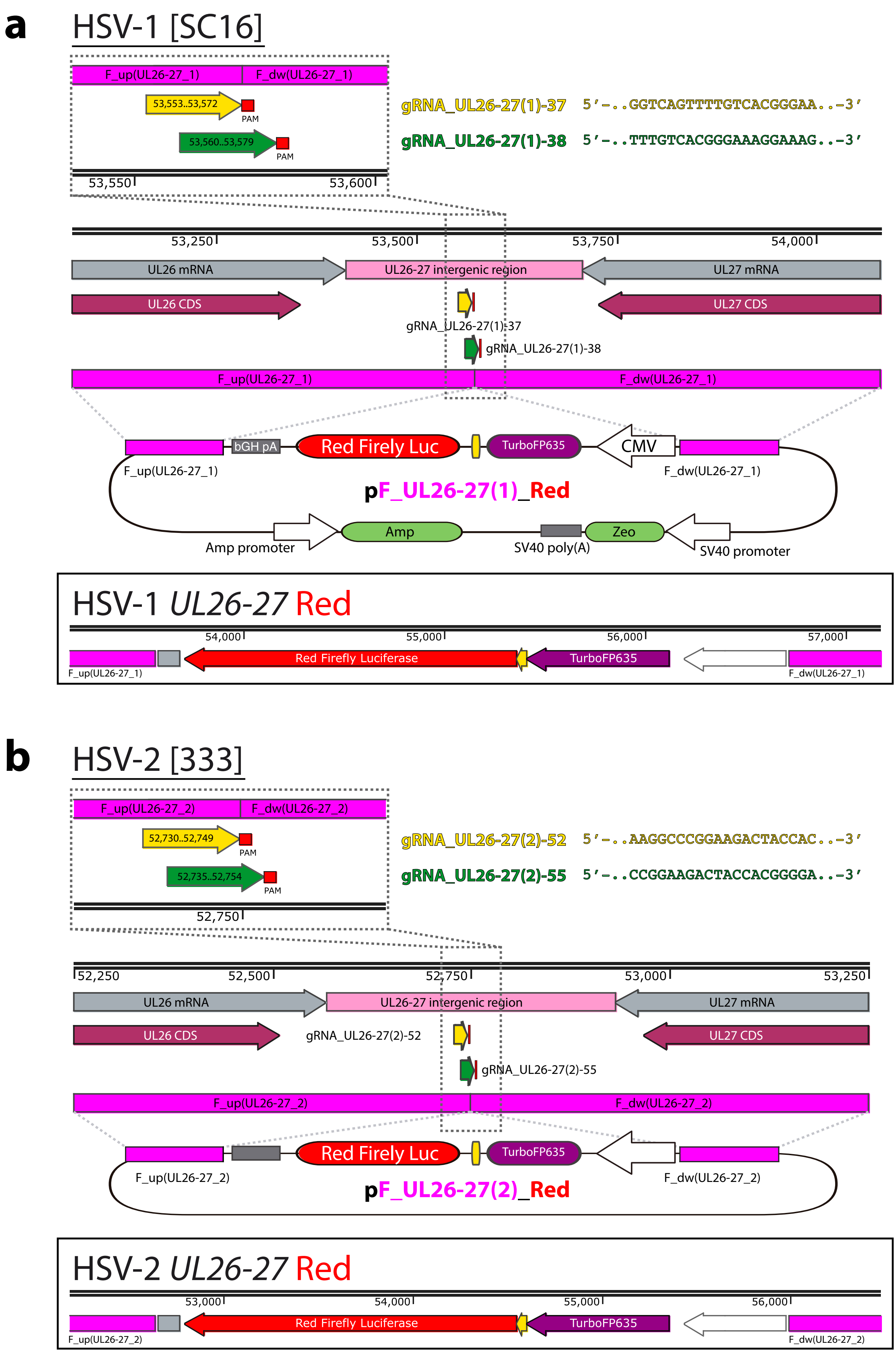


**Fig. S2. Schematic of recombinants HSV-1/2 *UL26-27* Red generation.** The intergenic region *UL26-27* in HSV-1 (**a**) and HSV-2 (**b**) genomes was simultaneously targeted at the indicated gRNA sites. By using the donor templates pF_UL26-27(1/2)_Red to induce HDR, HSV-1/2 *UL26-27* Red were obtained.

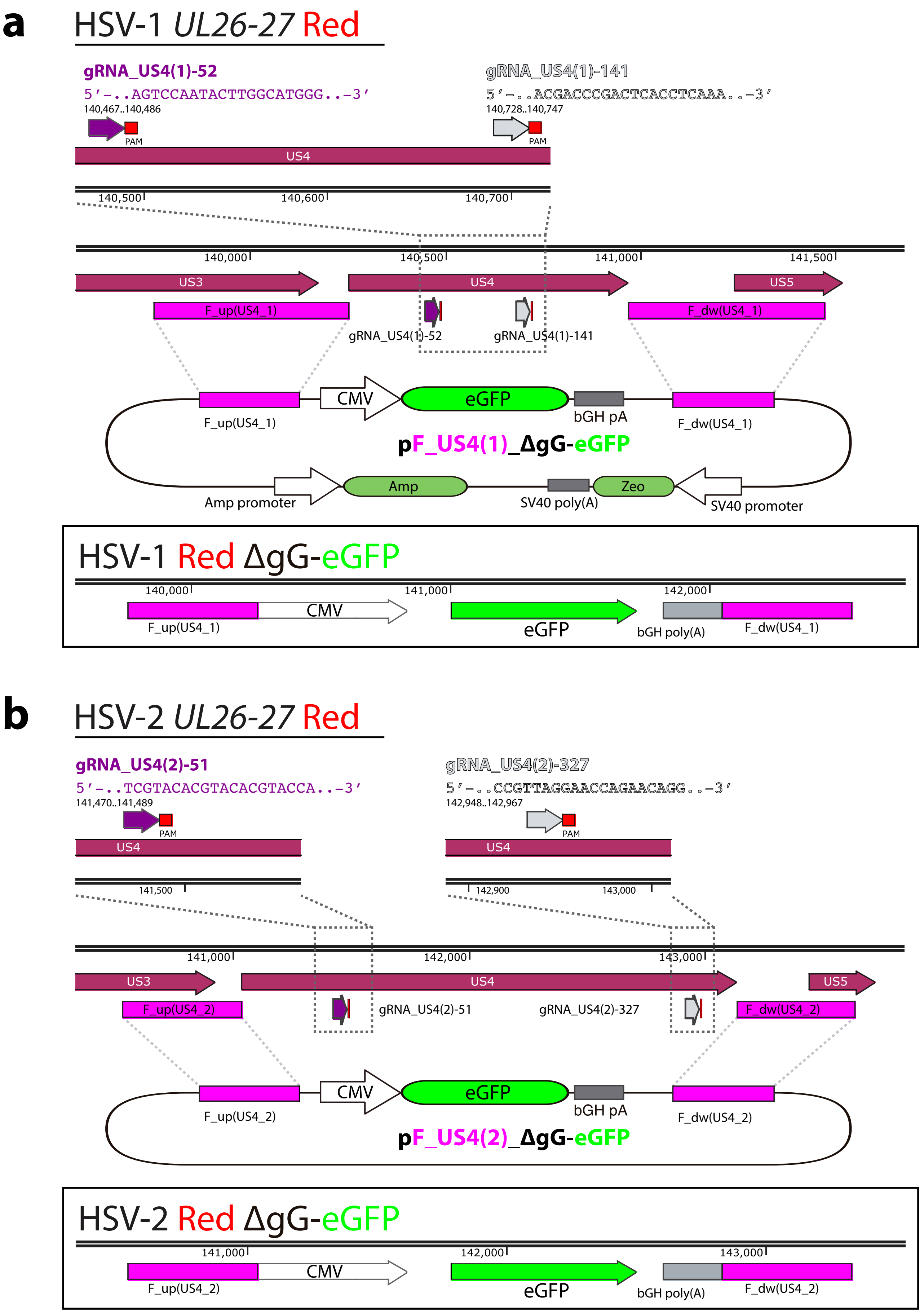


**Fig. S3. Schematic of recombinants HSV-1/2 Red ΔgG-eGFP generation.** The *US4* locus in HSV-1 (**a**) and HSV-2 (**b**) *UL26-27* Red genomes was simultaneously targeted at the indicated gRNA sites. HDR templates pF_US4(1/2)_ΔgG-eGFP were used to produce HSV-1/2 Red ΔgG-eGFP. The eGFP cassette completely replaced the *US4* gene in the mutant viral genomes.

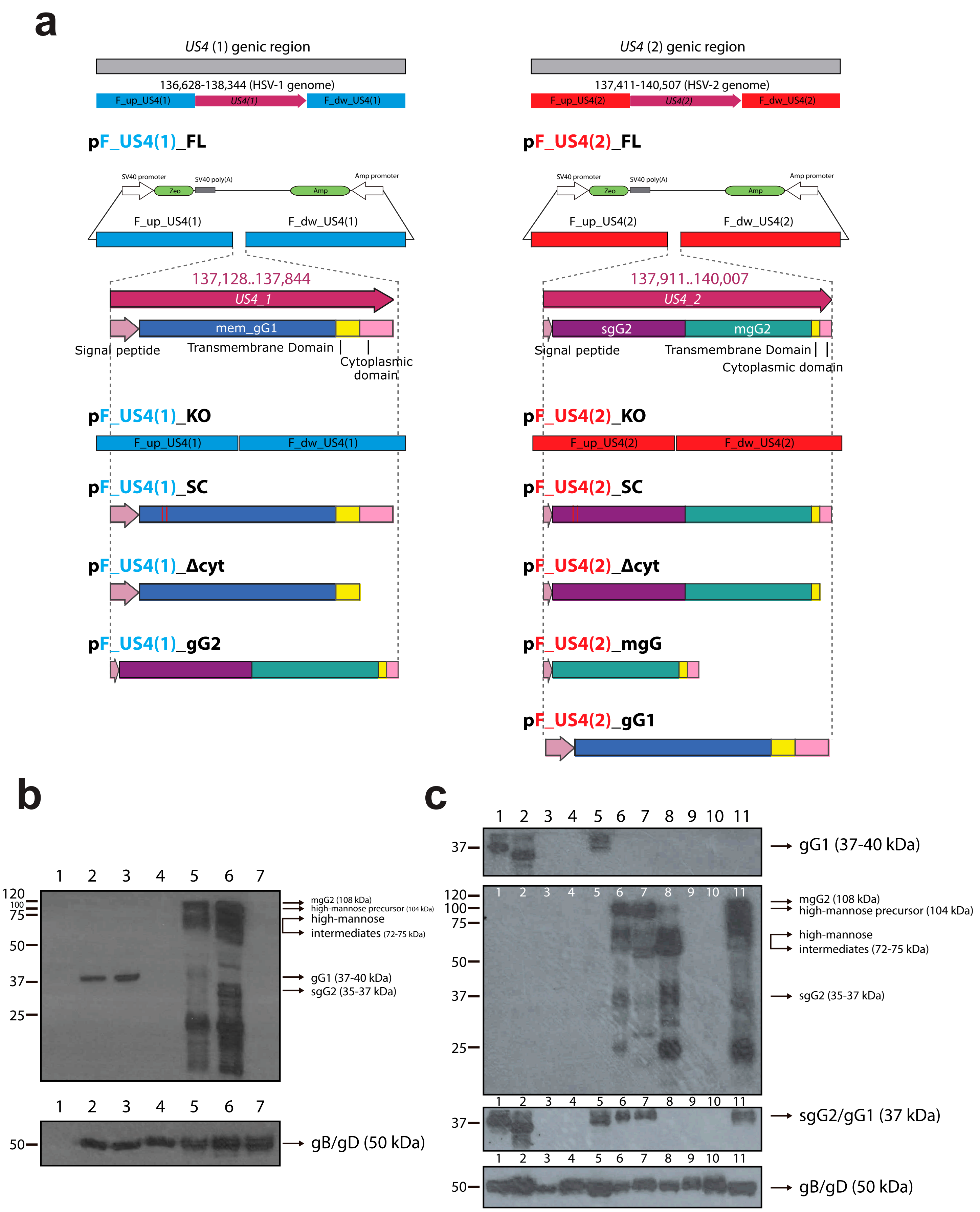


**Fig. S4. Donor plasmids used as HDR templates for the generation of a collection of mutants HSV-1/2 in the *US4* locus, and characterization of their gG expression.** (**a**) Schematic of the *US4* genic region in the wild-type HSV-1/2 genomes and modified plasmids used as HDR templates for each *US4* mutant virus. (**b, c**) Characterization of gG expression of *US4* mutant viruses by western blotting. Vero cells were infected at a MOI of 5 PFU/cell and harvested after 48 hpi. Western blot analysis was performed on cell extracts using anti-gG1 (Virus Corporation #H1379), anti-gG2 (Virus Corporation #1206), anti-sgG2/gG1 (polyclonal antibody in-house generated), and anti-gB/gD (control, gift from E. Tabares, Universidad Autonoma de Madrid). Vero cells in (**b**) were mock-infected (1) or infected with WT HSV-1 (2), HSV-1 Red (3), HSV-1 Red ΔgG-eGFP (4), WT HSV-2 (5), HSV-2 Red (6) and HSV-2 Red ΔgG-eGFP (7). Vero cells in (**c**) were infected with HSV-1 Red (1), HSV-1 Red gG-Δcyt (2), HSV-1 Red gG-KO (3), HSV-1 Red gG-SC (4), HSV-2 Red gG1 (5), HSV-2 Red (6), HSV-2 Red gG-Δcyt (7), HSV-2 Red gG-mgG (8), HSV-2 Red gG-KO (9), HSV-2 Red gG-SC (10) and HSV-1 Red gG2 (11). The position of the respective proteins is indicated by arrows. Molecular size markers in kDa are shown on the left. Only one viral clone out of two tested is shown.

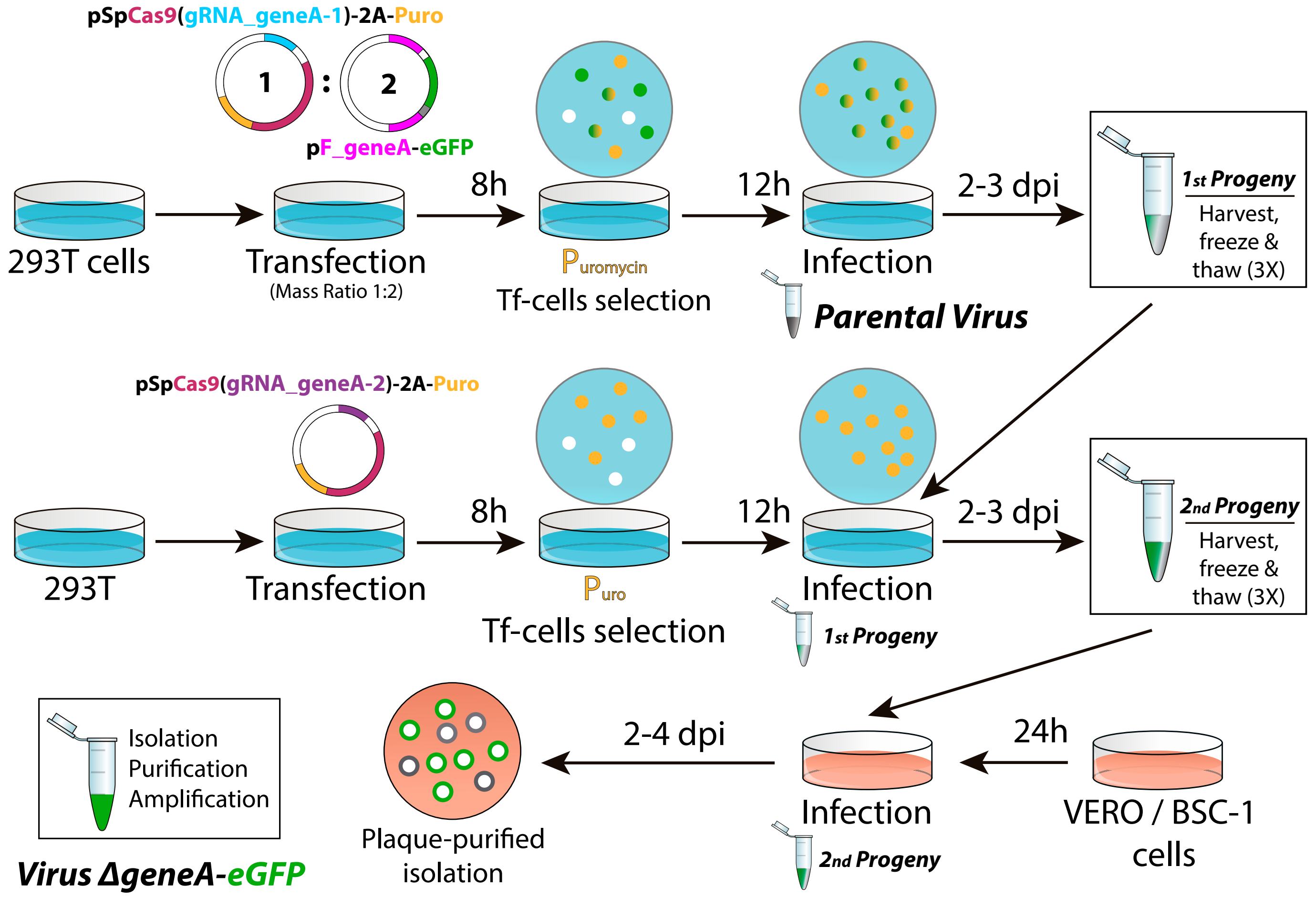


**Fig. S5. Flowchart of the Two-Progenies method applied to generate a large collection of recombinant viruses for a desired locus.** After selecting a gene of interest in the viral genome (e.g. gene *A*), 293T cells are transfected with the multicistronic pSpCas9-gRNA-1 vector (ΔNLS for cytoplasmic-replicating viruses), and the HDR template pF_geneA-eGFP. Then, cells are puromycin-selected and infected with the parental virus. Simultaneously, fresh 293T cells are transfected only with pSpCas9-gRNA-2 against gene *A*. After drug selection, cells expressing Cas9 and gRNA-2 are infected with the first viral progeny previously harvested. The second viral progeny is harvested and used to infect cells for identification and isolation of recombinant eGFP(+) plaques.

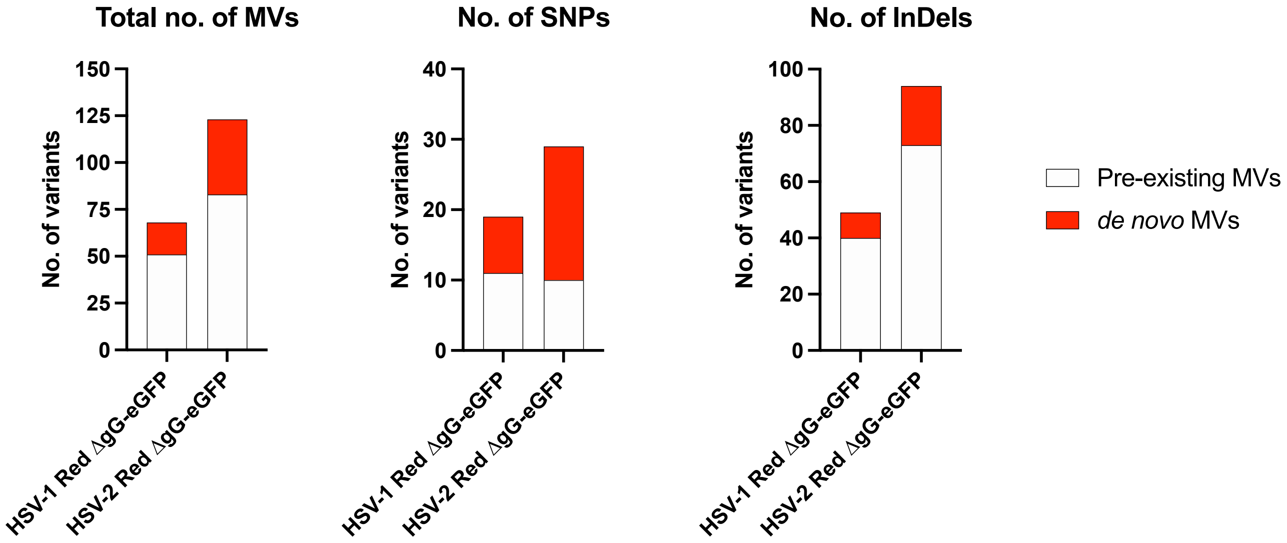


**Fig. S6. Genetic variant analysis of parental Cas9-derived HSV-1/2 Red ΔgG-eGFP.** Number of MVs, SNPs and InDels are plotted according to genetic variant analysis data for each parental recombinant virus. Histograms differentiate between pre-existing MVs observed in the corresponding parental virus (white), *de novo* appeared MVs (red).

### SUPPLEMENTARY METHODS

#### Generation of recombinant ECTV *195P*-eGFP

**Genomic insertion point of the VACV p7.5 promoter-eGFP cassette.**

This cassette was inserted into the *195P* locus of ECTV Naval genome replacing genomic coordinates 187,462 – 187,573.

**VACV p7.5 promoter-eGFP cassette sequence. 903 bp (5' – 3'):**

atcagtacccttcacctaattccaaacccacccgctttttatagtaagtttttcacccataaataataaatacaata
attaatttctcgtaaaagtagaaaatatattctaatttattgcacggtaaggaagtagatcataacaccatggtga
gcaagggcgaggagctgttcaccggggtggtgcccatcctggtcgagctggacggcgacgtaaacggcca
caagttcagcgtgtccggcgagggcgagggcgatgccacctacggcaagctgaccctgaagttcatctgca
ccaccggcaagctgcccgtgccctggcccaccctcgtgaccaccctgacctacggcgtgcagtgcttcagcc
gctaccccgaccacatgaagcagcacgacttcttcaagtccgccatgcccgaaggctacgtccaggagcgc
accatcttcttcaaggacgacggcaactacaagacccgcgccgaggtgaagttcgagggcgacaccctggt
gaaccgcatcgagctgaagggcatcgacttcaaggaggacggcaacatcctggggcacaagctggagta
caactacaacagccacaacgtctatatcatggccgacaagcagaagaacggcatcaaggtgaacttcaag
atccgccacaacatcgaggacggcagcgtgcagctcgccgaccactaccagcagaacacccccatcggc
gacggccccgtgctgctgcccgacaaccactacctgagcacccagtccgccctgagcaaagaccccaacg
agaagcgcgatcacatggtcctgctggagttcgtgaccgccgccgggatcactctcggcatggacgagctgt
acaagtaaaagggtcaagacaattctgcagatatcattaacaaac

Generation of donor plasmids for recombinant HSV-1/2 in the *US4* locus (see *"Plasmids"* section in Methods and **Fig. S4a**)

Genomic coordinates cloned into each plasmid correspond to HSV-1 strain SC16 partial genome (KX946970) and HSV-2 strain 333 complete genome (LS480640):

- pF_US4(1)_FL: 136,628 – 138,344 (KX946970)
- pF_US4(2)_FL: 137,411 – 140,507 (LS480640)
- pF_US4(1)_KO: 137,128 – 137,844 (deleted) (KX946970)
- pF_US4(2)_KO: 137,911 – 140,007 (deleted) (LS480640)
- pF_US4(1)_SC: 137,275 bp (deleted) and 137,281 – 137,282 ("T" insertion) (KX946970)
- pF_US4(2)_SC: 138,060 bp (deleted) and 138,066 (T → G) (LS480640)
- pF_US4(1)_ ∆cyt: 137,758 – 137,841 (KX946970)
- pF_US4(2)_ ∆cyt: 139,921 – 140,004 (LS480640)
- pF_US4(2)_mgG: 137,977 – 138,939 (LS480640)
- pF_US4(1)_ gG2: 137,128 – 137,844 (LS480640)
- pF_US4(2)_ gG1: 137,911 – 140,007 (KX946970)

Generation of recombinant HSV-1/2 *UL26-27* Red

**Plasmids**

To generate the donor vectors pF_UL26-27(1)_Red and pF_UL26-27(2)_Red (**Fig. S1a**), firstly, the HSV-1 *UL26-27* intergenic region (53,071 - 54,078; accession # KX946970), was amplified by PCR with primers F_up_UL26-27(1)_Fw and F_dw_UL26-27(1)_Rv. Then, it was cloned into the modified version of pcDNA3.1/Zeo(-) by the In-Fusion cloning system (Takara), obtaining pF_UL26-27(1). Similarly, the HSV-2 *UL26-27* intergenic region (52,250 - 53,249; accession # LS480640) was amplified with primers F_up_UL26-27(2)_Fw and F_dw_UL26-27(2)_Rv and cloned, producing pF_UL26-27(2).

To produce the Red cassette, the humanized far-red fluorescent protein TurboFP635 (scientific name *Katushka*) was amplified from pTurboFP635-N plasmid (Evrogen) with primers TurboFP635_Fw and TurboFP635_Rv. The 2A peptide (T2A) from *Thosea asigna* virus capsid protein was amplified from pSpCas9(BB)-2A-Puro V2.0 (Addgene #62988) with primers T2A_Fw and T2A_Rv. The pCMV-Red-Firefly-Luc vector was linearized by amplification with primers pCMV-RedLuc_Fw and pCMV-RedLuc_Rv, and then, together with TurboFP635 and T2A fragments, were In-Fusion cloned to construct pCMV-TurboFP635-T2A-RedFireflyLuc-bGH.

The CMV-TurboFP635-T2A-RedFireflyLuc-bGH fragment (red cassette) was amplified with primers CMV-Red(UL26-27_1)_Fw and CMV-Red(UL26-27_1)_Rv, and then, In-Fusion cloned in reverse orientation into linearized pF_UL26-27(1), giving pF_UL26-27(1)_Red. Similarly, primers CMV-Red(UL26-27_2)_Fw and CMV-Red(UL26-27_2)_Rv were used to amplify the red cassette, being In-Fusion cloned into linearized pF_UL26-27(2) to produce pF_UL26-

27(2)_Red. pF_UL26-27(1) was linearized with primers F_dw_UL26-27(1)_Fw and F_up_UL26-27(1)_Rv, while pF_UL26-27(2) with primers F_dw_UL26-27(2)_Fw and F_up_UL26-27(2)_Rv.

gRNAs targeting the *UL26-27* intergenic region were designed with Protospacer Workbench software, based on ROI 53,412-53,706 for HSV-1 and 52,568-52,932 for HSV-2 (Table S10). gRNA_UL26-27(1)-37 and #-38 were selected based on their location at the F_up/dw(UL26-27(1)) border (Fig. S2a), as well as gRNA_UL26-27(2)-52 and -55 regarding HSV-2 (Fig. S2b), ensuring the disruption of the complementary sequence for each gRNA in recombinant HSV *UL26-27* Red viruses. Primers gRNA_UL26-27(1)-37_Fw and gRNA_UL26-27(1)-37_Rv were annealed and cloned to generate plasmid pSpCas9(gRNA_UL26-27(1)-37). Likewise, primers gRNA_UL26-27(1)-38_Fw and gRNA_UL26-27(1)-38_Rv, obtaining pSpCas9(gRNA_UL26-27(1)-38); primers gRNA_UL26-27(2)-52_Fw and gRNA_UL26-27(2)-52_Rv to produce pSpCas9(gRNA_UL26-27(2)-52), and primers gRNA_UL26-27(2)-55_Fw and gRNA_UL26-27(2)-55_Rv to obtain pSpCas9(gRNA_UL26-27(2)-55).

**One-step generation by CRISPR/Cas9**

293T cell monolayers with 3 x $10^5$ cells/well in 6-well plates were transfected simultaneously with 1 µg of donor vector pF_UL26-27(1)_Red, 1 µg of pSpCas9(gRNA_UL26-27(1)-37) and 1 µg of pSpCas9(gRNA_UL26-27(1)-38). After puromycin selection, cells were infected with WT HSV-1 (MOI of 1) to generate HSV-1 *UL26-27* Red. Similarly, HSV-2 *UL26-27* Red was produced by simultaneous transfection with pF_UL26-27(2)_Red, pSpCas9(gRNA_UL26-27(2)-52) and pSpCas9(gRNA_UL26-27(2)-55), and then, puromycin selection and infection with WT HSV-2 (MOI of 1). After 2 dpi, cells were harvested, subjected to three freeze-thaw cycles and 10-fold serially diluted to infect Vero cells. HSV-1/2 *UL26-27* Red were identified by fluorescent microscopy, plaque-purified and characterized.

**Genomic insertion point of the red cassette**

The red cassette was inserted into the HSV-1 *UL26-27* intergenic region (53,572 – 53,573), and into the HSV-2 genome (52,749 – 52,750).

**Red cassette sequence, bGH-RedFireflyLuc-T2A-TurboFP635-CMV.**

**3126 bp (5’ – 3’):**

gatgcaatttcctcattttattaggaaaggacagtgggagtggcaccttccagggtcaaggaaggcacgggg
gaggggcaaacaacagatggctggcaactagaaggcacagtcgaggctgattttgcggccgctcatcacat
cttggccacgggtttcttcaggatctcccggatggccctgccgtcgatcttgccggtcaggccctttggcacttcgt
ccacgaatctcacgccgcctctcagccgcttggcgttggacacctggctggcgacgtagtccatcacttctttctc
ggtcatgttcttgccggattccagcaccaccacggcgccaggcagctcgccggccacaggatctggcacccc
ggccacgccggcgtcgaagatgctggggtgctgcagcaggacgctttccagctcggcagggggcacctgat
agcccttgtacttgatcaggctcttcagccggtccacgatgaagaagtgcttctcttcgtcgtagtagccgatgtc
gccggtgtgcagccagccctcttcgtcgatcagctctttggtggcctcggggttgttcacgtagcccttcatcagc
atggggcccttcacgcacacttcgccccgtctgttggggcccaggctcttcttggtgtccaggtcgatcactttgg
ccttgaacaggggcaccaccttgccgctggctccaggcttgtcgtcgccctcgggggtgatgatgatggcgct
ggttgtctcggtcaggccgtagccctgccgcacgccgggcagattgaaccgcctggcgacggcctctcccac
ttctttgctcagggggggctccgccgctggcgatctccaccaggttgctcaggtcgtacttgttcagcagctcgctct
tgttcaggatggcgaacagggtgggcaccagaatcacgtaggtgcacttgtagtcctgcagggttttcaggaa
ggtttcctcgtcgaacttggtcagcatcaccacccggaagccgcagatcaggtagcccagggtggtgaacat

gccgaagccgtggtggaagggcaccacggtcagcacggcggtgccgggggacacctggttgccgtagatg

gggtccctggcgtggctgaaccgggtcacggtgttctcgtgggtcagctgcacgcccttgggcaggccggtgc

tgccgctgctgttcatgatcagggccacctgttctttccggtccacctccacggtcttgaagctgctggcctggaa

gccaggggggggtgttccgcttgatgaaggtgtccaggcactggtagccccggtagtccaccttgctgtccagg

atcacgatggtcttgatggtggtcacggttttctgcacggtgatgactttgtccaggcccttcttgctgctgaacacg

atggtgggcttgctgatgcccaggctgtgcaccagctcccgcagggtgtagatctcgttggtgggagccacgc

ccacgccgatgaacaggccggcgatcacggggatgaagaattcctcgcagttctcgctgcacagggcgatc

cggccgtccaccaccaggccgtagttctgcagagccttgcccaggcagcagctcttttccaggtactcggcgt

agctgtagtccacgccggtcacggcgttggtgaaggcaatggcgcccagcttggcgtatctttccatgtacttcc

gcagctgggtgccggcgctgccttcctcgatggggtagaagggcttggggcccaccacgatgttctcgtcgtttt

ccatattttccattgggccaggattctcctcgacgtcaccgcatgttagcagacttcctctgccctcgctgtgcccc

agtttgctaggcaggtcgcagtacctggccacagccatctcgtgctgctcgacgtaggtctctttgtcggcctcctt

gattctttccagtctcctgtccacgaagtagaagccgggcatcttgaggttcttagcgggtttcttggatctgtatgt

ggtcttgagggagcagtgcaggtagcccccgcccacgagcttcagggccatctggctatggcctctcaggcc

gctgtcagcggggtacagcatctcggtgctggcctcccagccgagtgttttcttctgcatcacagggccgttgga

tgggaagttcaccccgttgatcttgacgttgtagatgaggcagccgttctggaggctggtgtcctgggtagcggt

cagcacgcccccgtcttcgtatgtggtgatcctctcccatgtgaagccctcagggaaggactgcttaaagaagt

cggggatgccctgggtgtggttgataaaggttttgctgccgtacatgaagctggtagccaggatgtcgaaggc

gaaggggagagggccgccctcgaccaccttgatcttcatggtctgggtgccctcgtagggcttgccttcgccct

cggatgtgcacttgaagtggtggtcgttcacggtgccctccatgtacagtttcatgtgcatgttctcggtgatcagc

acgctatcctcacccaccatggtggcggatcccctatagtgagtcgtattaatttcgataagccagtaagcagtg

ggttctctagttagccagagagctctgcttatatagacctcccaccgtacacgcctaccgcccatttgcgtcaatg

gggcggagttgttacgacattttggaaagtcccgttgattttggtgccaaaacaaactcccattgacgtcaatgg

ggtggagacttggaaatccccgtgagtcaaaccgctatccacgcccattgatgtactgccaaaaccgcatca

ccatggtaatagcgatgactaatacgtagatgtactgccaagtaggaaagtcccataaggtcatgtactgggc

ataatgccaggcgggccatttaccgtcattgacgtcaatagggggcgtacttggcatatgatacacttgatgtac
tgccaagtgggcagtttaccgtaaatactccacccattgacgtcaatggaaagtccctattggcgttactatggg
aacatacgtcattattgacgtcaatgggcgggggtcgttgggcggtcagccaggcgggccatttaccgtaagtt
atgtaacg

Generation of recombinant HSV-1/2 Red ΔgG-eGFP

**Plasmids**

The CMV-eGFP-bGH fragment (eGFP cassette) was HF amplified from plasmid pEGFP-N3 (Takara) with primers EGFP(US4-1)_Fw and EGFP(US4-1)_Rv, or primers EGFP(US4-2)_Fw and EGFP(US4-2)_Rv, and In-Fusion cloned in forward orientation into previously linearized pF_US4(1)/(2) backbones, giving pF_US4(1)/(2)_ΔgG-eGFP, respectively (**Fig. S1b**). pF_US4(1) was linearized from pF_US4(1)_FL with primers F_dw_US4(1)_Fw and F_up_US4(1)_Rv, while pF_US4(2) from pF_US4(2)_FL with primers F_dw_US4(2)_Fw and F_up_US4(2)_Rv.

gRNAs targeting the *US4* CDS were based on ROI 137,128 - 137,844 for HSV-1 (accession # KX946970), and 137,911 - 138,910 (forward) / 139,008 - 140,007 (reverse) for HSV-2 (accession # LS480640) (**Fig. S3**, Table S11). Primers gRNA_US4(1)-52_Fw and gRNA_US4(1)-52_Rv were annealed and cloned to generate plasmid pSpCas9(gRNA_US4(1)-52); primers gRNA_US4(1)-141_Fw and gRNA_US4(1)-141_Rv to make pSpCas9(gRNA_US4(1)-141); primers gRNA_US4(2)-51_Fw and gRNA_US4(2)-51_Rv to obtain pSpCas9(gRNA_US4(2)-51) and primers gRNA_US4(2)-327_Fw and gRNA_US4(2)-327_Rv to produce pSpCas9(gRNA_US4(2)-327).

##### One-step generation by CRISPR/Cas9

293T cell monolayers with 3 x $10^5$ cells/well in 6-well plates were transfected simultaneously with 1 µg of donor vector pF_US4(1)_ΔgG-eGFP, 1 µg of pSpCas9(gRNA_US4(1)-52) and 1 µg of pSpCas9(gRNA_US4(1)-141). After puromycin selection, cells were infected with HSV-1 *UL26-27* Red (MOI of 1) to generate HSV-1 Red ΔgG-eGFP. Similarly, HSV-2 Red ΔgG-eGFP (Fig. S5b) was produced by transfection with pF_US4(2)_ΔgG-eGFP, pSpCas9(gRNA_US4(2)-51) and pSpCas9(gRNA_US4(2)-327), and then, puromycin selection and infection with HSV-2 *UL26-27* Red (MOI of 1). After 2 dpi, cells were harvested, subjected to three freeze-thaw cycles and 10-fold serially diluted to infect Vero cells. Recombinant HSVs were identified by fluorescent microscopy, plaque-purified and characterized.

##### Genomic insertion point of the *eGFP* cassette.

The *eGFP* cassette was inserted into the HSV-1 genome between 137,127 – 137,845 reference positions, replacing the *US4* gene (137,128-137,844). Similarly, it was inserted into the HSV-2 genome between 137,910 – 140,008 coordinates, also replacing the *US4* gene (137,911 - 140,007).

##### eGFP cassette sequence, CMV-eGFP-bGH (CMV and bGH selected regions for gRNA design are highlighted in bold, italic and underline).

**1795 bp (5' – 3'):**

***<u>gacattgattattgactagttattaatagtaatcaattacggggtcattagttcatagcccatatatggagtt</u>***

***<u>ccg</u>***cgttacataacttacggtaaatggcccgcctggctgaccgcccaacgacccccgcccattgacgtcaat aatgacgtatgttcccatagtaacgccaatagggactttccattgacgtcaatgggtggactatttacggtaaact

gcccacttggcagtacatcaagtgtatcatatgccaagtacgccccctattgacgtcaatgacggtaaatggcc

cgcctggcattatgcccagtacatgaccttatgggactttcctacttggcagtacatctacgtattagtcatcgctat

taccatggtgatgcggttttggcagtacatcaatgggcgtggatagcggtttgactcacggggatttccaagtctc

caccccattgacgtcaatgggagtttgttttggcaccaaaatcaacgggactttccaaaatgtcgtaacaactcc

gccccattgacgcaaatgggcggtaggcgtgtacggtgggaggtctatataagcagagctctctggctaacta

gagaacccactgcttactggcttatcgaaattaatacgactcactatagggagacccaagctggctagcgcta

ccggactcagatctcgagctcaagcttcgaattctgcagtcgacggtaccgcgggcccgggatccaccggtc

gccaccatggtgagcaagggcgaggagctgttcaccggggtggtgcccatcctggtcgagctggacggcga

cgtaaacggccacaagttcagcgtgtccggcgagggcgagggcgatgccacctacggcaagctgaccctg

aagttcatctgcaccaccggcaagctgcccgtgccctggcccaccctcgtgaccaccctgacctacggcgtg

cagtgcttcagccgctaccccgaccacatgaagcagcacgacttcttcaagtccgccatgcccgaaggctac

gtccaggagcgcaccatcttcttcaaggacgacggcaactacaagacccgcgccgaggtgaagttcgagg

gcgacaccctggtgaaccgcatcgagctgaagggcatcgacttcaaggaggacggcaacatcctggggca

caagctggagtacaactacaacagccacaacgtctatatcatggccgacaagcagaagaacggcatcaag

gtgaacttcaagatccgccacaacatcgaggacggcagcgtgcagctcgccgaccactaccagcagaaca

cccccatcggcgacggccccgtgctgctgcccgacaaccactacctgagcacccagtccgccctgagcaaa

gaccccaacgagaagcgcgatcacatggtcctgctggagttcgtgaccgccgccgggatcactctcggcatg

gacgagctgtacaagtaaagcggccgccactgtgctggatatctgcagaattccaccacactggactagtgg

atccgagctcggtaccaagcttaagtttaaaccgctgatcagcctcgactgtgccttctagttgccagccatctgt

tgtttgcccctcccccgtgccttccttgaccctggaaggtgccactcccactgtcctttcctaataaaatgaggaa

attgcatc***gcattgtctgagtaggtgtcattctattctggggggtggggtggggcaggacagcaaggg***

***ggaggattgggaagacaatagcaggcatgctggggatgcggtgggctctatgg***
